# Non-Toxic Virucidal Macromolecules Show High Efficacy Against Influenza Virus Ex Vivo and In Vivo

**DOI:** 10.1101/2020.03.18.996678

**Authors:** Ozgun Kocabiyik, Valeria Cagno, Paulo Jacob Silva, Yong Zhu, Laura Sedano, Yoshita Bhide, Joelle Mettier, Chiara Medaglia, Bruno Da Costa, Samuel Constant, Song Huang, Laurent Kaiser, Wouter L. J. Hinrichs, Anke Huckeriede, Ronan Le Goffic, Caroline Tapparel, Francesco Stellacci

## Abstract

Influenza is one of the most widespread viral infections worldwide and represents a major public health problem. The risk that one of the next pandemics is caused by an influenza strain is very high. It is very important to develop broad-spectrum influenza antivirals to be ready for any possible vaccine shortcomings. Anti-influenza drugs are available but they are far from ideal. Arguably, an ideal antiviral should target conserved viral domains and be virucidal, *i*.*e*. irreversibly inhibit viral infectivity. Here, we describe a new class of broad-spectrum anti-influenza macromolecules that meets these criteria and displays exceedingly low toxicity. These compounds are based on a cyclodextrin core modified on its primary face with long hydrophobic linkers terminated in 6’sialyl-N-acetyllactosamine (6’SLN) or 3’SLN. SLN enables nanomolar inhibition of the viruses while the hydrophobic linkers confer irreversibility to the inhibition. The combination of these two properties allows for efficacy in vitro against several human or avian influenza strains, as well as against a 2009 pandemic influenza strain ex vivo. Importantly, we show that, in mice, the compounds provide therapeutic efficacy when administered 24h post-infection allowing 90% survival as opposed to no survival for the placebo and oseltamivir..

---

Influenza viruses are among the most infective viruses.^[1,2]^ Every year different influenza strains infect a large fraction of both the animal and human population^[3]^ endangering infants, the elderly and immunocompromised people, which are at high risk of hospitalization and death, due to influenza-related complications.^[4–8]^ As a result, seasonal influenza has yearly a remarkable socio-economic impact. Respiratory diseases can cost a significant fraction of the total health expenditures in developed and mainly in developing countries.^[9,10]^ Because influenza mutates so rapidly, the development of a lifelong vaccine is still a major challenge.^[11–13]^ Vaccine development would pose even higher challenges when we focus on the occasional pandemics instead of yearly outbreaks. In such a case, the development time of a new vaccine would represent a serious risk. Furthermore, even in the presence of a vaccine, reaching reasonable vaccination coverage is far from a foregone conclusion. As a consequence, the risk of a new pandemic, such as the Spanish-flu, is still present and recognised as one of the top threats to global health.^[14–16]^

Naturally, the second line of defense after vaccines, are antiviral drugs. A number of anti-influenza drugs are currently approved: neuraminidase inhibitors such as zanamivir and oseltamivir, ion channel inhibitors such as amantadine, fusion inhibitors such as umifenovir (only in Russia and China) and polymerase inhibitor such as baloxavir marboxil, that was recently approved in US and Japan. Yet, it is recognized that the efficacy of current drugs is far from ideal. Concerns about these drugs range from significant side effects to the appearance of drug-resistant viruses after a short period of use.^[17]^ Given the importance of this issue, a number of other antivirals are in clinical trials.^[18–25]^ The majority of these drugs are monoclonal antibodies^[26,27]^ that inhibit the fusion of the virus to the host-cell. Although they are promising, it is likely that they will be considerably costly due to their manufacturing processes. Furthermore, monoclonal antibodies are expected to be good prophylactic drugs, but their efficacy in therapeutic administration (*i*.*e*. post infection) is a matter of intense research.

An ideal anti-influenza drug should be broad-spectrum, by targeting a highly conserved part of the virus and, in order to avoid loss of efficacy due to the dilution in body fluids, have an irreversible effect, *i*.*e*. be virucidal. Obviously, this drug needs to be truly non-toxic. There are quite a few research lines on the development of molecules that target conserved parts of the virus.^[28–40]^ To the best of our knowledge, these compounds are all reversible in their action and hence could face major hurdles when translating into drugs. To the best of our knowledge no compound with broad-spectrum anti-influenza efficacy has shown convincing post-infection results in vivo of the type that we present here.

The search for virucidal (*i*.*e*. irreversible) drugs with limited toxicity has been very challenging. Polymers bearing hydrophobic groups have previously been reported to show virucidal activity or enhanced antiviral activity.^[41,42]^ We have also shown that gold nanoparticles^[43]^ and β-cyclodextrins^[44]^ modified with 11-undecane sulfonic acid display a virucidal mechanism against a wide range of heparan sulfate proteoglycans (HSPGs) binding viruses, with no cellular toxicity. These compounds were capable of exerting forces that ultimately deform the virus particle. The chemical structure of the ligand was shown to be essential in order to achieve such irreversible inhibition.

Here, we adopted a similar strategy in order to target human influenza viruses and avian influenza viruses. 6’ sialyl-N-acetyllactosamine (6’SLN) and 3’ sialyl-N-acetyllactosamine (3’SLN), that bind to hemagglutinin (HA) trimers of influenza strains with high affinity^[45]^, were grafted onto the primary face of β-cyclodextrins through a series of different linkers. The structure of all modified cyclodextrins discussed in this work is shown in **Figure 1**. Of note, since each HA trimer has three sialic acid binding pockets, we aimed to modify the β-CDs with three trisaccharides. Therefore, the cyclodextrins are not fully modified but they all bear, on average, a comparable number of trisaccharides (Figure S1 and S2), determined using ^1^H Nuclear Magnetic Resonance Spectroscopy (NMR). In vitro dose-response assays against influenza A/Netherlands/602/2009 (H1N1) strain (A/NL/09), were conducted to compare the inhibitory activity of these molecules (Figure 1 and S3). The infection was quantified with immunocytochemical assays at 24 hours post-infection (hpi). β-CDs bearing a sufficiently long, hydrophobic linker and 6’SLN end-group (**C6-6’, C11-6’**, and **C14-6’**) showed strong inhibitory activity against infection with influenza A/NL/09, having EC50 values in the nanomolar range. On the other hand, the β-CD with a shorter linker, **C1-6’**, poorly protected from the infection. Introducing a sufficiently long linker clearly enhanced the end-group flexibility; hence the inhibitory concentrations decreased. The EC50 was comparable (yet slightly higher) when the hydrophobic linker was replaced with a hydrophilic PEG8 linker (**P8-6’**).

**Figure 1:**
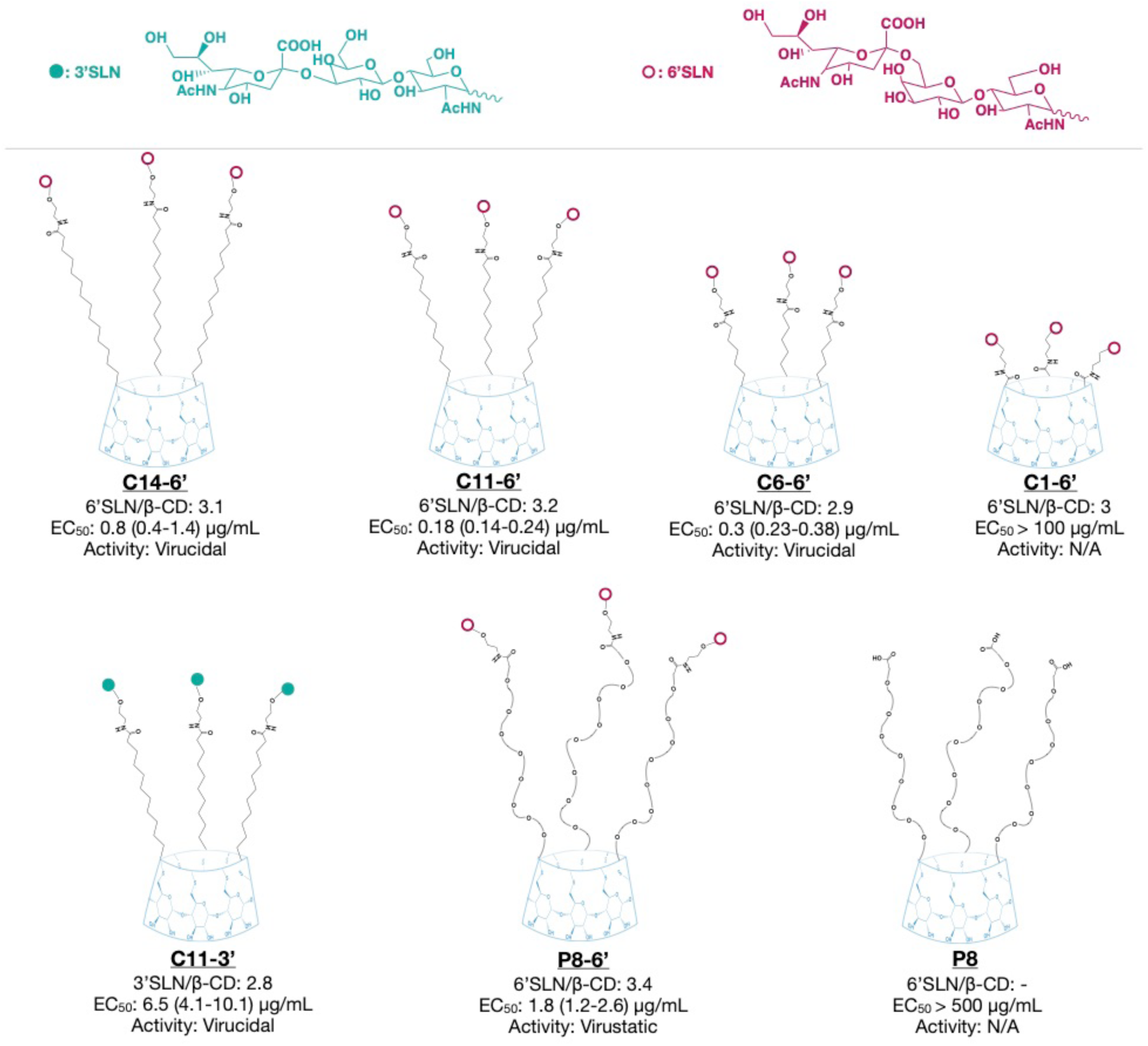
Summary of the modified cyclodextrins. The average number of 6’SLN or 3’SLN per β-CD was calculated by ^1^H NMR. The representative chemical structures of modified cyclodextrins were constructed based on NMR results. EC_50_ represents the half-effective concentrations on MDCK cells at 24 hpi against A/NL/09 with the respective 95% confidence interval (CI) (Figure S3). N/A: not assessable.

In order to demonstrate viral inhibition is due to trisaccharide group but not due to β-CD or the linker bearing carboxylic acid end group, we also tested β-CD solely modified with PEG8 linker **(P8)**. P8 did not inhibit influenza A/NL/09, even at very high material concentrations.

**C11-6’**, the molecule that showed the best inhibitory activity against A/NL/09, displayed strong antiviral activity against human influenza strains from both the A (H1N1 and H3N2) and the B type (Table 1 and Figure S4). Importantly, it inhibited very recent A (H1N1) and B clinical strains (from the 2017/2018 influenza season), isolated from patients in the University Hospital of Geneva and passaged only once in MDCK cells. **C11-6’** did not show any antiviral activity against HSV-2, an HSPG-binding virus, indicating specificity of the compound for sialic acid dependent viruses.

**Table 1:**
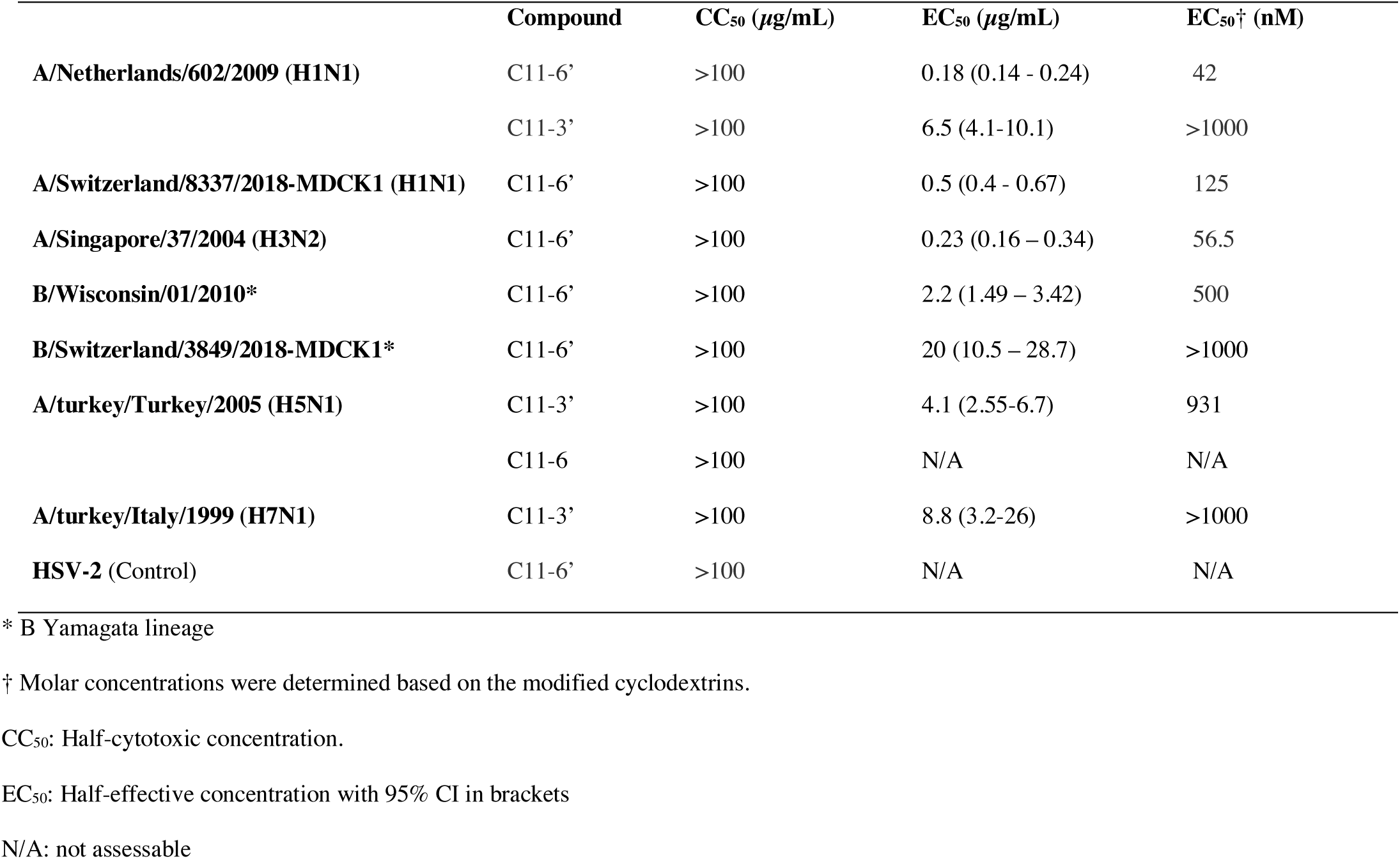
Inhibitory activity of C11-6’ and C11-3’ against different influenza strains.

6’SLN is known to be specific to human influenza strains, whereas 3’SLN is preferred by avian influenza strains as a primary attachment point.^[46]^ To prove the generality of our approach, especially against influenza strains that are known to have the ability of crossing the species barrier, we synthesized and tested **C11-3’** (Figure 1) against avian influenza strains. We show that **C11-3’** inhibits both H5N1 and H7N1 avian strains, at 4.1 and 8.8 μg/ml concentrations respectively (see Table 1). *De facto*, these results confirm the antiviral strategy adopted against human strains. We additionally tested whether **C11-3’** could inhibit the human strain A/NL/09 and whether **C11-6’** would also be active against the avian strain, H5N1. **C11-3’** displayed a good inhibitory activity against A/NL/09 (Figure 1 and table 1), whereas **C11-6’** did not show any activity against H5N1 (table 1 and Figure S5). These results are in line with previous literature comparing the binding affinities of avian and human strains to the different types of sialic acids^[46,47]^. Avian influenza strains (particularly H5N1 strains) preferentially bind to alpha -2,3 linked sialic acid, which has a thin and straight trans conformation. On the other hand, the wider sialic acid binding site of human strains can accommodate both the bulky cis conformation of alpha -2,6 linked sialic acid and the narrower -2,3 linked sialic acid^[46,47]^.

The synthesis of similar compounds sharing the β-cyclodextrin core and the 6’SLN moiety but different linkers allowed us to highlight a structural feature conferring irreversible inhibitory activity i.e. virucidal action (**Figure 2** and S6). This feature differs from the one described in our previous work where only the length of the linkers was compared^[43]^. Here, we hypothesized that one of the key components of an irreversible viral inhibition is that the binding moiety (here 6’SLN) is borne by a sufficiently long hydrophobic linker. A hydrophilic linker such as PEG should not be capable of generating forces that permanently inactivate the virus. To test this hypothesis, we compared **C11-6’** and **P8-6’**. These two compounds differ solely in the hydrophobicity of the linker and show comparable inhibitory activity against A/NL/09, as shown on the left in figure **2a** and **2b**. Virucidal assays were conducted as previously described^[43]^ to compare the mechanism of inhibition of these compounds, *i*.*e*., virucidal (irreversible) or virustatic (reversible). Briefly, amounts of the compounds that provide complete protection (10 μg of **C11-6’** and 50 μg of **P8-6’**) were incubated with the virus for 1h. Serial dilutions of the inocula were conducted followed by evaluation of the infectivity. In the case of **P8-6’**, the right graph in **2a** shows that while at the initial concentration complete protection was present, upon dilution the difference with the infectivity of the control sample (virus alone) was lost, *i*.*e*. the inhibitory effect was found to be reversible (virustatic). In the case of **C11-6’** the right graph in **2b** shows that complete protection was kept upon dilution and the graphs in **2c** show that this property was conserved against a number of different strains.

**Figure 2:**
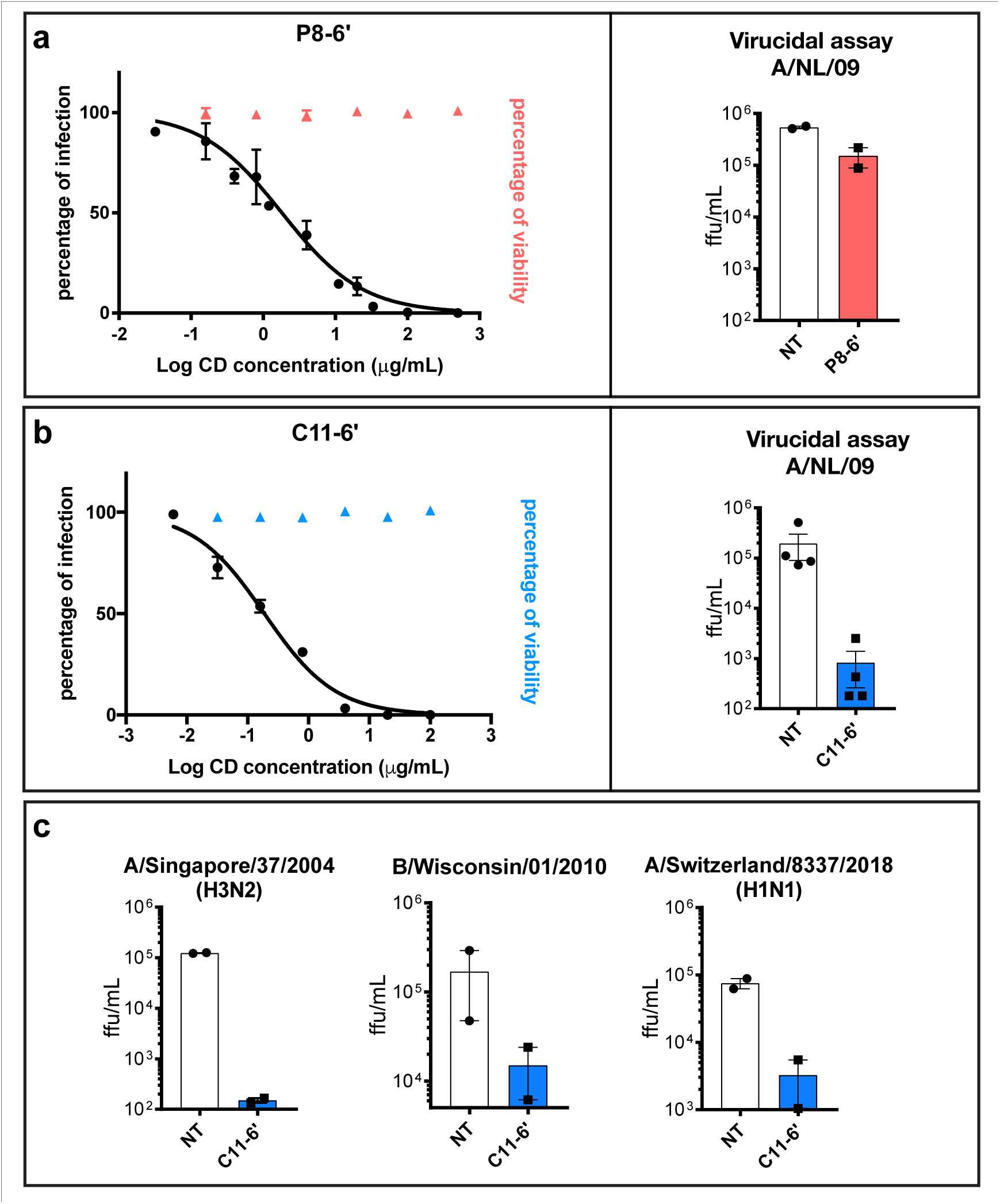
Antiviral activity comparison of C11-6’ and P8-6’ in vitro. Panels (**a**) and (**b**) show on the graphs on the left the inhibitory activity of each compound against A/NL/09, superimposed with the results of the cell viability assays. Both of the compounds inhibit the virus in the dose-response assay. In the virucidal assays on the right, **C11-6’** reduced the virus titer by 1000 times, whereas the infection was fully recovered in the case of **P8-6’**. Hence **C11-6’** has an irreversible inhibitory effect on the virus while the effect of **P8-6’** is reversible. Virucidal activity of **C11-6’** against other influenza strains was further investigated confirming its irreversible activity independently of the strain (**c**). Note that in the figure’s axes **ffu** stands for focus forming units and **NT** for non-treated. In (**c**) the following viral strains were tested: A/Singapore/37/2004 (H3N2), B/Wisconsin/01/2010 and A/Switzerland/8337/2018-MDCK1 (H1N1) (**c**). Results are mean and SEM of 2 independent experiments performed in duplicate.

To further compare **C11-6’** and **P8-6’**, we performed ex vivo experiments in MucilAir**®**, a 3D model of human airway reconstituted epithelia. These air-liquid interface cultures perfectly mimic both the pseudostratified architecture (basal, ciliated and goblet cells) and the barrier defence mechanism (i.e. the mucociliary clearance and epithelial cell immunity) of the human upper respiratory epithelium, the main site of influenza virus replication in humans. Ex vivo experiments were conducted with clinical H1N1 pandemic 09 strain (A/Switzerland/3076/16) that has not been passaged in cells to exclude any adaptation bias. **C11-6’** or **P8-6’** (50 µg/tissue) and the virus (10^4^ RNA copies/tissue) were first added simultaneously on the apical surface of the tissues, without prior incubation. After four hours, the inocula were removed, the tissues were washed and the progress of the infection was monitored on a daily basis with qPCR performed on viruses isolated from the apical washes of the tissue, without any re-addition of the molecule. **C11-6’** completely prevented virus replication throughout the entire course of the experiment, while **P8-6’** slightly reduced viral replication the first two days post-infection (dpi) but not thereafter (**Figure 3a**).

**Figure 3:**
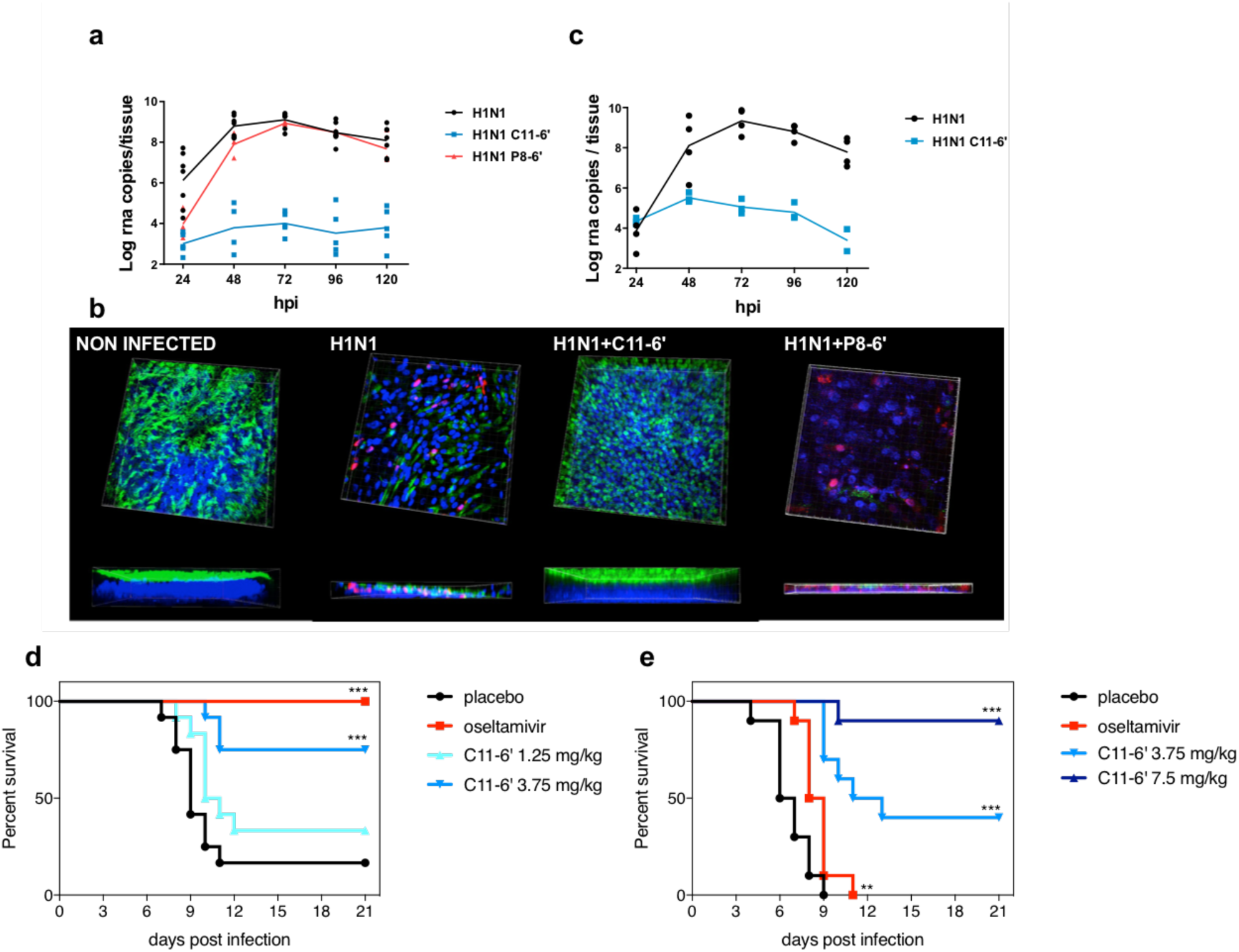
Ex vivo (a to c) and in vivo (d to f) inhibitory activity of C11-6’. **Ex vivo, C11-6’** provided a full protection against clinical H1N1 pandemic 09 strain in co-treatment condition, whereas P8-6’ only provided a minor protection in the beginning of the infection (**a**) Immunofluorescence at 7 days post-infection (co-treatment condition) confirms the protection provided by **C11-6’**. Red: monoclonal antibody influenza A, blue: DAPI, green: β-IV-tubulin (marker of ciliated cells). The thickness of each tissue is shown at the bottom of the corresponding image (**b**). **C11-6’** also showed high efficacy in post-treatment condition (**c**). Results of (a) and (c) are mean and SEM of 2 to 4 independent experiments with intra-experimental duplicates. Images of (b) are representative of 10 images taken for each condition. **In vivo**, mice (12/group) were intra-nasally treated with PBS or **C11-6’** or by oral gavage with oseltamivir and infected with A/California/09. Subsequent treatments were administered daily for the following 5 days, image shows the survival curve (**d**). Mice were infected and treated with **C11-6’** or oseltamivir at 24 hpi and daily for the 4 following days, images show the survival curves (**e**) ***p<0.001, **p<0.01

Moreover, in the tissues treated with **C11-6’**, the inhibition of viral replication was also reflected by the absence of infected cells and the undisturbed morphology of the treated tissues, strikingly different from the untreated or **P8-6’**-treated tissues (Figure 3b). Immunofluorescence images and the lack of lactate dehydrogenase (LDH) release in the apical washes demonstrated that the ciliated cell layer as well as the physiological cilia beating and tissue integrity were preserved (Figure 3b and S7). In stark contrast, the untreated tissue or the **P8-6’**-treated controls, presented reduced thickness due to alteration of the ciliated cell layer, and presence of infected cells (Figure 3b). To exclude that the residual viral level detected by qPCR in the treated tissues was related to active replication, we kept the tissues in culture for 23 days but never observed an increase in viral titer over time, while the untreated tissues were persistently shedding virus as previously reported^[48]^ (Figure S8). Importantly, ex vivo experiments were conducted also in more stringent post-treatment conditions in which **C11-6’** (30 µg/tissue) was administrated every 24 h and for 4 days, starting at 1 dpi to mimic a therapeutic administration. Also in these conditions, the compound showed a remarkable inhibitory activity, proving its potential as a therapeutic agent (Figure 3c). In the same ex vivo model we also evaluated the biocompatibility of high doses of **C11-6’**, administered daily. **C11-6’** did neither show any cytotoxic nor pro-inflammatory activity in the above described conditions (Figure S9).

Lastly, in vivo experiments were conducted with BALB/c mice administered first with **C11-6’** (1.25 or 3.75 mg/kg) and immediately after with A/California/09 via the intranasal route and subsequently treated daily for 5 days. The weights of the mice were measured daily in order to estimate the impact of **C11-6’** administration on the infected-animal’s physiological condition (weight loss variation is shown in Figure S10). Significant increase in survival was observed in presence of **C11-6’** 3.75 mg/kg (9/12 mice) and the oseltamivir 30 mg/kg/day (10/10 mice) if compared to placebo control (3/12 mice). Additional experiments were carried out with administration of 1.25 mg/kg of **C11-6’** at the time of infection by measuring the viral load in the lungs and the viral titer in the broncho-alveolar lavages at 2dpi (Figure S11). The significative reduction in presence of 1.25 mg/kg of **C11-6’** demonstrates that a single administration before infection is able to significantly decrease the infectious titer of the virus. Subsequently we performed post-treatment experiment, in which mice were treated at 8 (Figure S12) or 24 hpi (Figure 3d) with **C11-6’** (3.75 mg/kg or 7.5 mg/kg) or oseltamivir (30 mg/kg/day). In presence of **C11-6’** with the start of treatment at 24hpi 9 out of 10 mice survived to the viral challenge in the 7.5 mg/kg group and 4 in the 3.75 mg/kg group, in contrast with oseltamivir in which group 0 out of 10 survived. Collectively, these results suggest a potent prophylactic and therapeutic capacity of the **C11-6’** compound in vivo.

In summary, we present here a new design rule to produce effective, non-toxic, virucidal compounds against influenza virus. To dissect the relationship between the antiviral mechanism of action and the structure of our newly designed virucidal we compare, for the first time, its efficacy with that of a highly similar compound displaying a virustatic activity. We show that if the inhibitory effect is reversible, i.e. virustatic, the antiviral efficacy is lost when moving from in vitro to ex vivo. On the other hand, the virucidal counterpart of the same molecule keeps its efficacy is maintained from in vitro all the way to in vivo. Importantly, we show in vivo results with remarkable effect on mice survival for a pandemic strain of H1N1 with the compound given intranasally 24h post-infection. Therefore, we believe that our approach to design non-toxic virucidal macromolecules has an outstanding potential for the prevention and the treatment of not only human, but also avian influenza infections.

## Experimental Section

Detailed information on the experimental procedures can be found in the Supporting Information.

## Acknowledgements

The work was supported by the Leenaards Foundation (grant 4390 to L.K., C.T. and F.S.) and the Swiss National Science Foundation (Sinergia grant CRSII5_180323 to F.S. and C.T.). The mouse model work was done at the Institute for Antiviral Research at Utah State University. Funding was provided by the National Institute of Health contract number HHSN272201700041I, Task Order A30, from the Respiratory Diseases Branch, Division of Microbiology and Infectious Diseases, National Institute of Allergy and Infectious Diseases, National Institutes of Health, USA.

## Author contributions

Conceptualization, O.K., V.C., C.T. and F.S.; Methodology, O.K., V.C. and R.L.G. Investigation, O.K. and Y.Z. synthetized the cyclodextrins; O.K., V.C., J.B., J.M. and C.M. performed the in vitro experiments; V.C. performed the ex vivo experiments; L.S., B.D.C. and R.L.G. performed the in vivo experiments. Writing – Original Draft, O.K., V.C., C.T. and F.S. Writing – Review & Editing, all authors. Funding Acquisition, C.T., L.K. and F.S.; Resources, S.H. and S.C. Supervision, W.L.J.H., A.H., R.L.G., C.T., F.S.

