## supplementary data for "Non-Toxic Virucidal Macromolecules Show High Efficacy Against Influenza Virus Ex Vivo and In Vivo"

### Supporting Information

### 1.Supplemantary figures

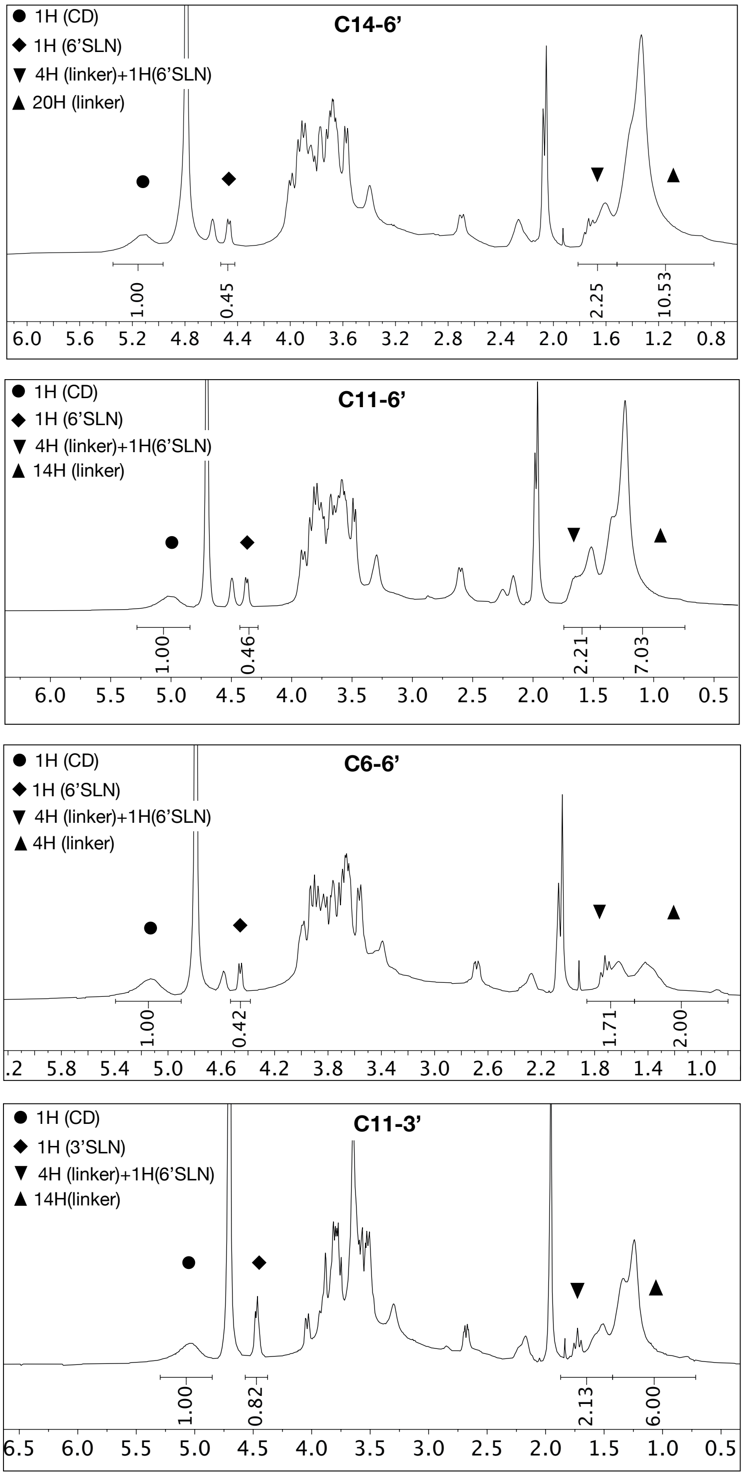


**Figure S1:** $^{1}$H NMR studies were conducted in order to characterize the modified cyclodextrins. The peaks corresponding to cyclodextrin, linker and trisaccharide were demonstrated with symbols. Note that the peak corresponding to cyclodextrin is the sum from 7 glucose subunits, each contributing 1H.

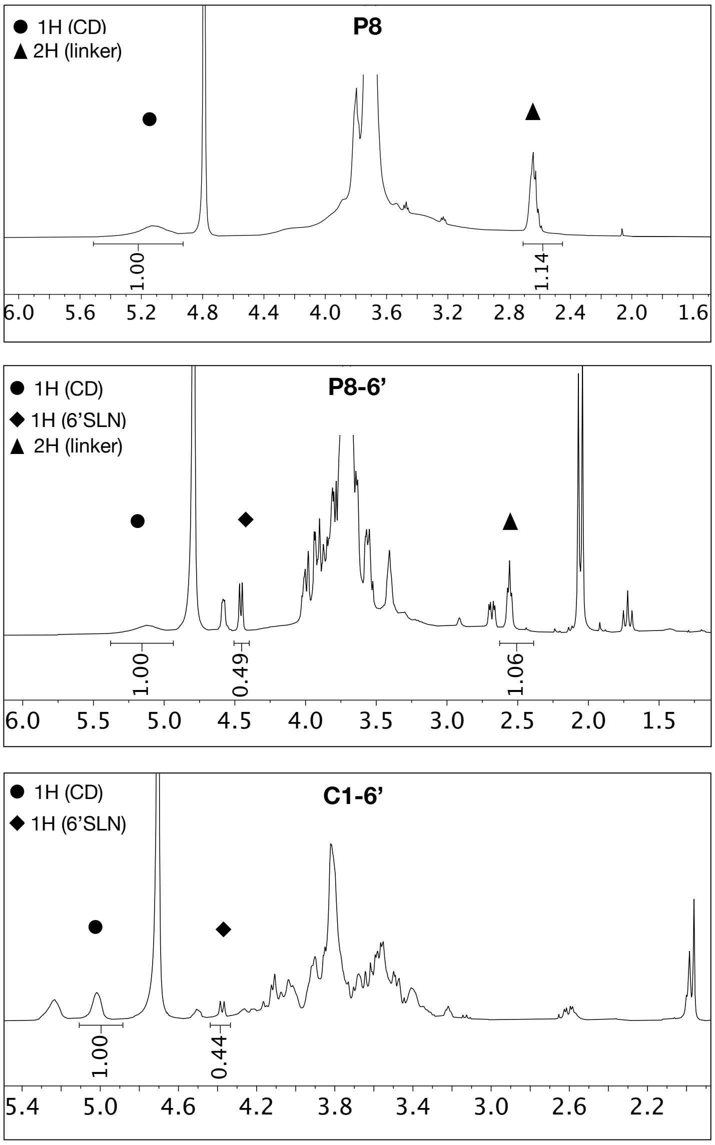


**Figure S1:** (Continued)

| | **Number of spacers/CD** | **Number of SLN/CD** | **Molecular weight (kDa)** |
|---|---|---|---|
| **C14-6'** | 3.5 | 3.1 | ~ 4.3 |
| **C11-6'** | 3.5 | 3.2 | ~ 4.3 |
| **C6-6'** | 3.4 | 2.9 | ~ 3.8 |
| **C11-3'** | 3.1 | 2.8 | ~ 3.9 |
| **P8** | 3.9 | - | ~ 3.2 |
| **P8-6'** | 3.7 | 3.4 | ~ 5.6 |
| **C1-6'** | - | 3 | ~ 3.6 |

**Figure S2:** Average number of 6'SLN or 3'SLN per ß-cyclodextrin determined by $^{1}$H NMR and molecular weights were estimated based on $^{1}$H NMR results.

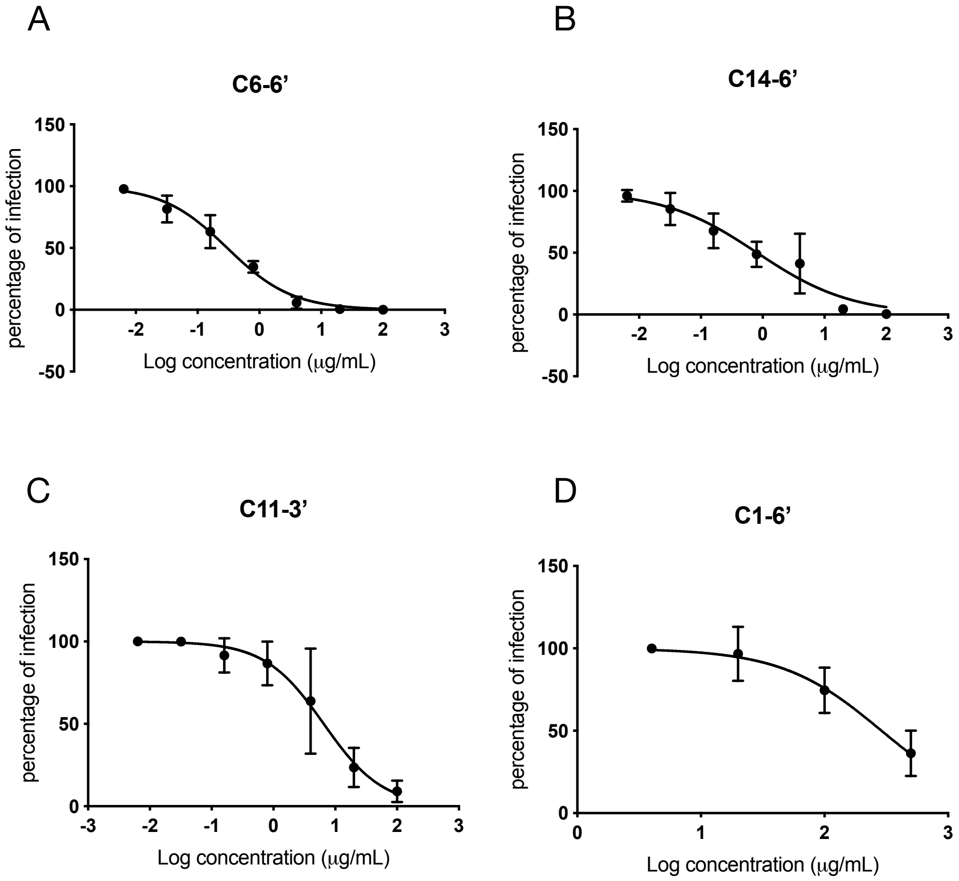


**Figure S3:** Dose-response curves demonstrating the antiviral activity of C6-6' (A), C14-6' (B), C11-3' (C) and C1-6' (D) against A/Netherlands/2009 H1N1.

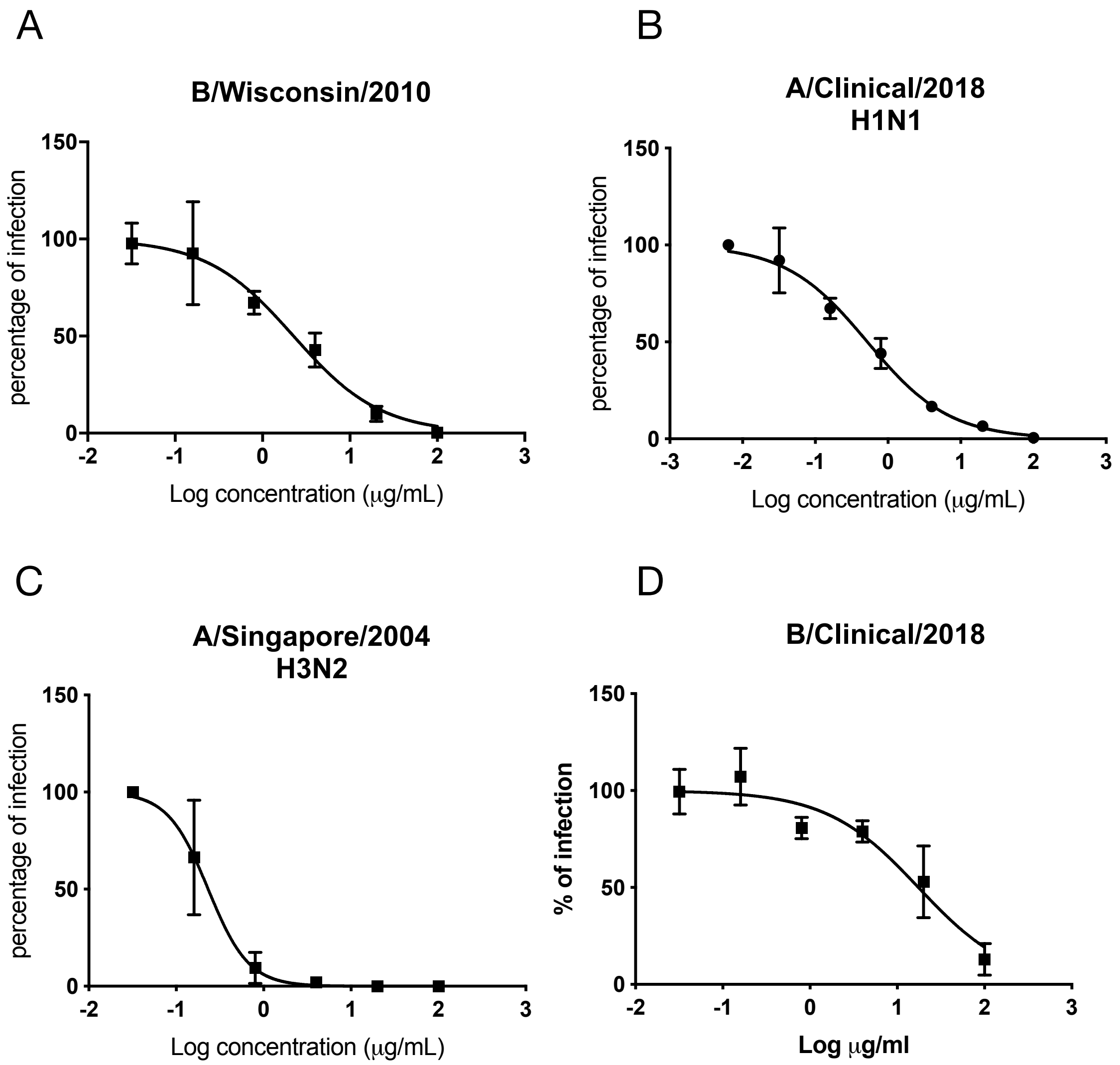


**Figure S4:** Dose-response curves demonstrating the antiviral activity of C11-6’ against B/Wisconsin/2010 (A), A/Clinical/2018 H1N1 (B), A/Singapore/2004 H3N2 (C) and B/Clinical/2018 (D)

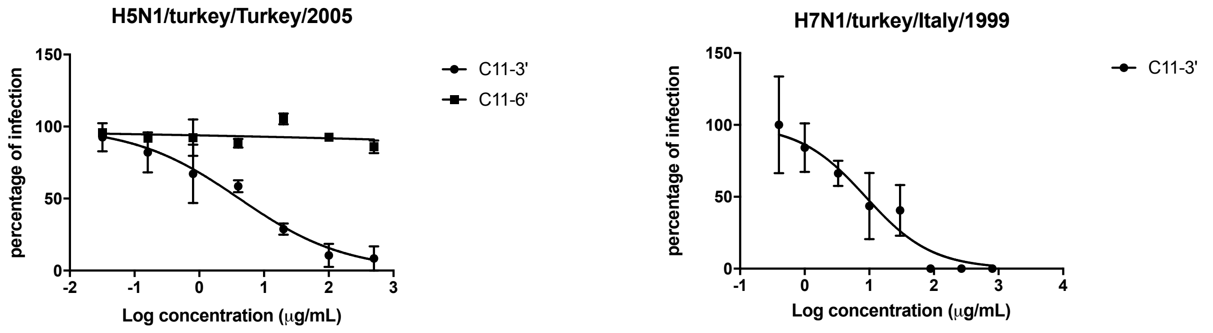


**Figure S5:** Dose-response curves demonstrating the antiviral activity of C11-3' against avian strains A/turkey/Turkey/2005 H5N1 (A) and A/Turkey/Italy /977/1999 H7N1 (B). A comparison experiment was also conducted with C11-6' in the case of A.

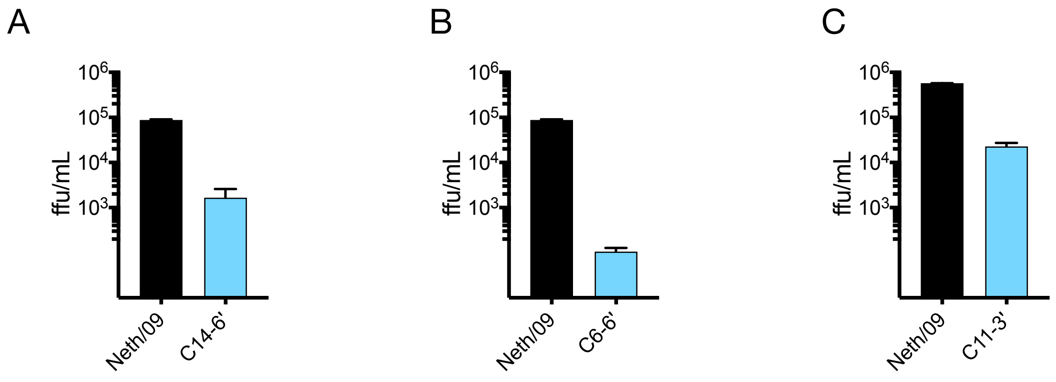


**Figure S6:** Virucidal activity of the C6-6' (A), C14-6' (B) and C11-3' (C) against A/Netherhands/2009 (H1N1). The experiments were performed with a compound concentration of 100 μg/mL.

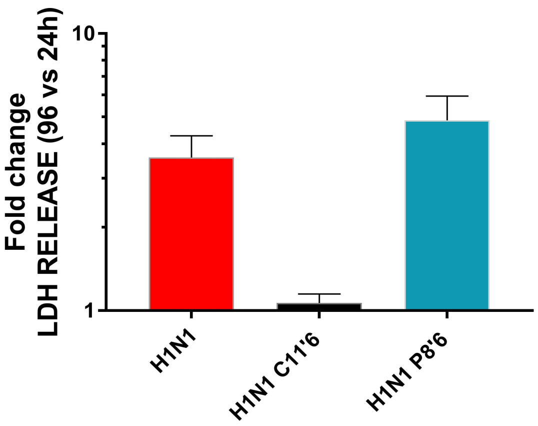


**Figure S7:** LDH release from infected tissues. Tissues were infected and treated with C11-6' (50 $\mu$g) or P8-6' at the time of infection. Apical washes performed at 96 and 24 hpi were subjected to LDH measurement. Results are mean and SEM of triplicates.

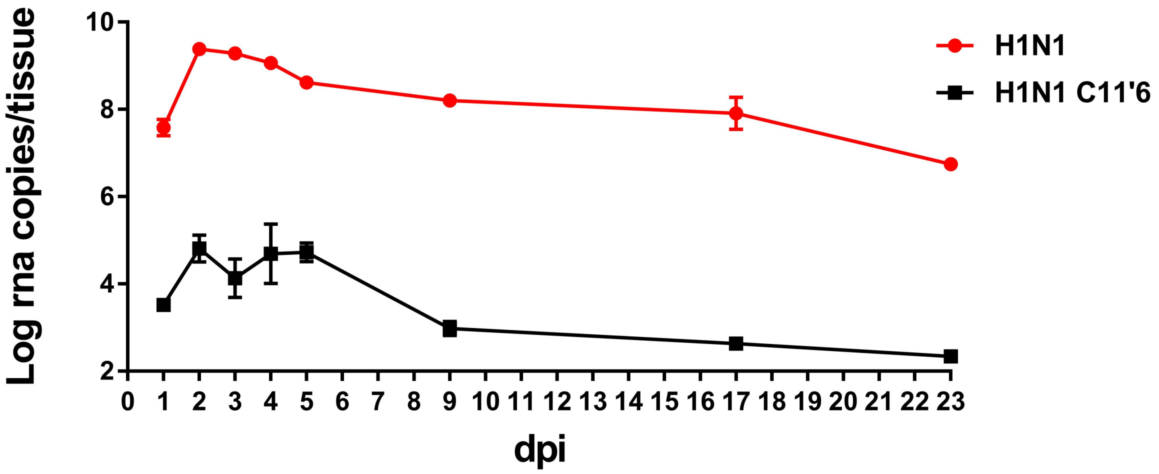


**Figure S8:** Long co-treatment experiment. Tissues were treated and infected with C11-6' (50 $\mu$g) at the time of infection. Daily apical washes were collected for the first 5 days, subsequently at 9, 17 and 23 days, with a wash the previous day in order to evaluate daily virus production. Results are mean and SD of a single experiment performed in duplicate.

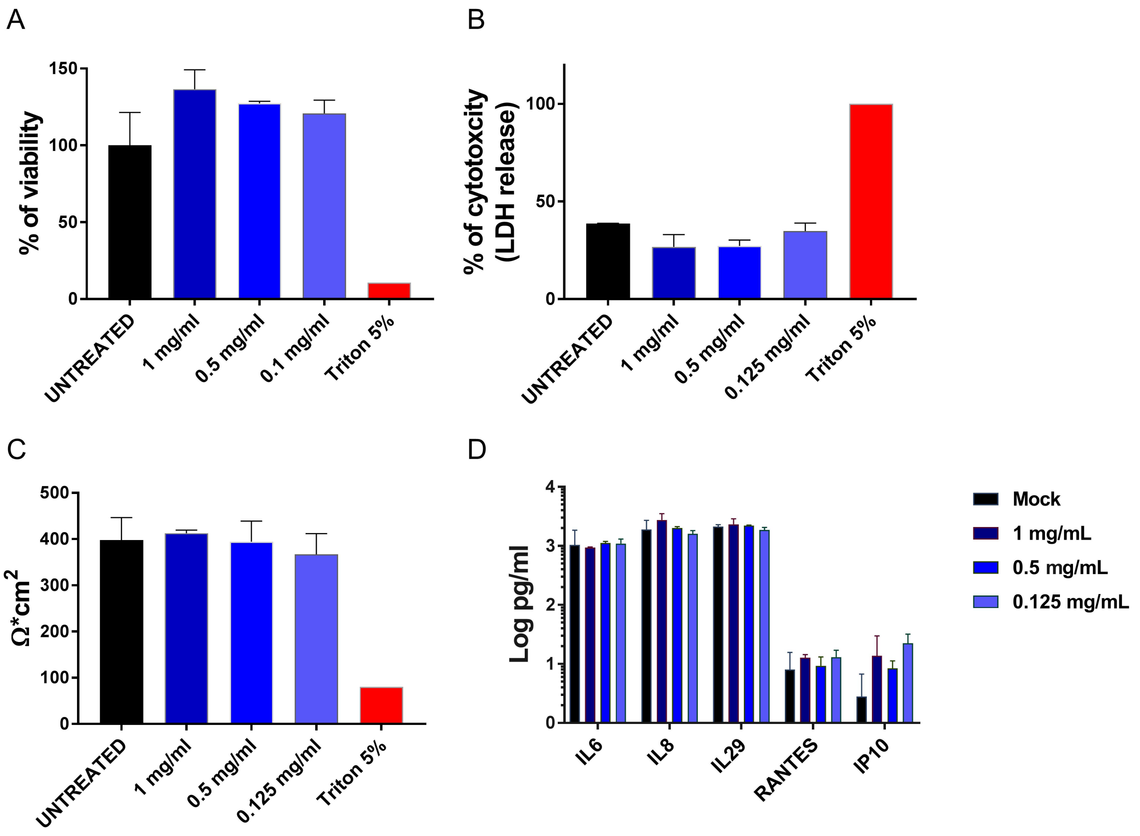


**Figure S9**: Ex vivo toxicity. Tissues were treated with different doses of C11'6 or equal volume of medium or triton 5% with daily addition. At 96 hours post treatment tissues were subjected to: A) MTT assay, B) LDH assay in which the viability of the tissues was evaluated, C) trans epithelial resistance evaluation and D) ELISAs assay to evaluate the release of pro-inflammatory cytokines. LDH and ELISA were performed on collected basal medium was collected and subjected to: B) LDH assay and D) The experiments are mean and SEM of 2 independent experiments performed in duplicates.

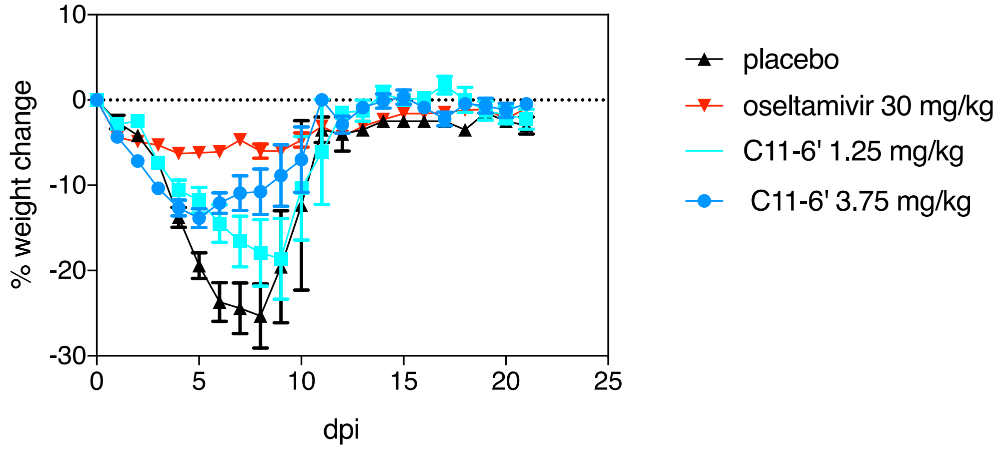


**Figure S10**. Effects of C11’6 treatments on body weights during the co-treatment test in BALB/c mice. Treatments were administered intranasally for 5 days. Data points represent the group mean and standard error of the percent change in weight relative to the first day of treatment.

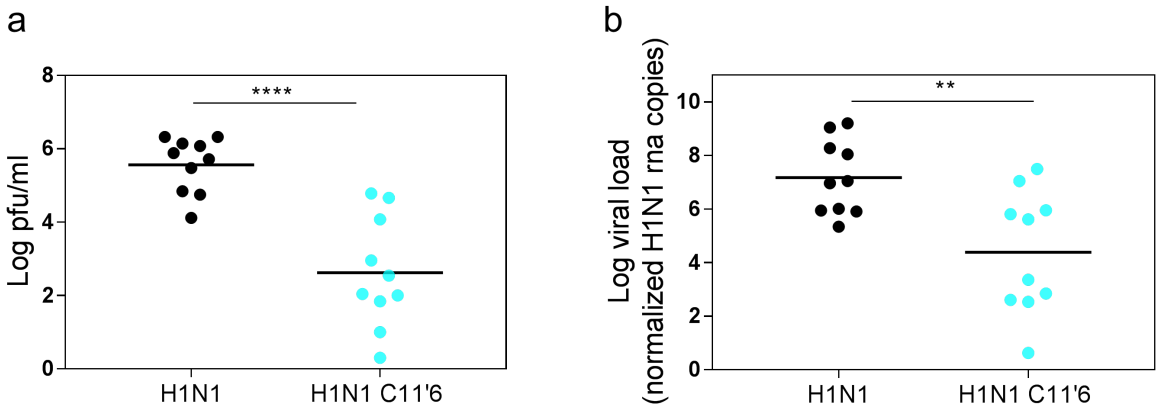


**Figure S11.** Effects of C11-6’ treatment on viral titer (a) and viral load (b) and of co-treated BALB/c mice. Treatment was administered intranasally once. Viral load (b) was calculated from homogenized lung homogenates while viral titer from bronchoalveolar lavage (b).

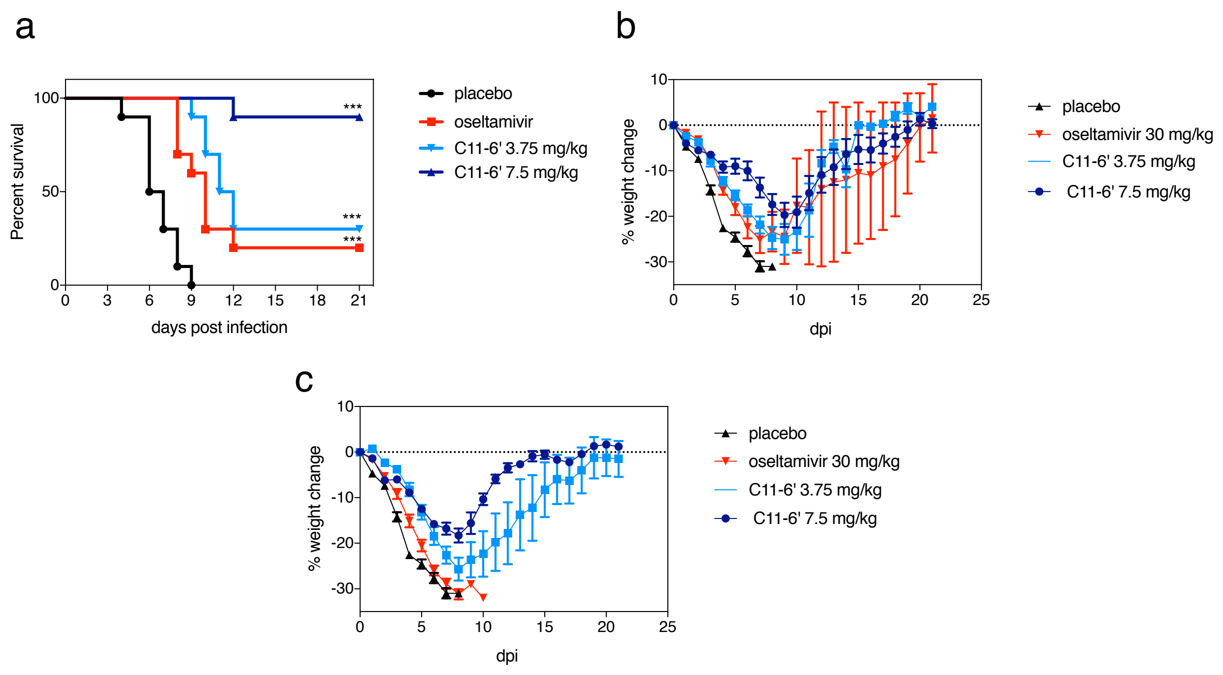


**Figure S12**. Effects of C11'6 treatments during the post-treatment test in BALB/c mice. Treatments were administered intranasally for 5 days starting at 8 hpi or 24 hpi. (a) Survival curves at 8hpi. (b) Weight change with start of treatment at 8hpi. (c) Weight change with start of treatment at 24hpi. Data points represent the group mean and standard error of the percent change in weight relative to the first day of treatment. ***$p<0.001$

### 2.Methods

#### Synthesis of Modified Cyclodextrins

**Chemicals:** Neu5Acα(2,6)-Galβ(1-4)-GlcNAc-β-ethylamine and Neu5Acα(2,3)-Galβ(1-4)-GlcNAc-β-ethylamine was purchased from TCI chemicals. Heptakis-(6-deoxy-6-mercapto)-beta-Cyclodextrin and carboxymethyl-beta-cyclodextrin sodium salt was purchased from Cyclodextrin-Shop. 11-dodecenoic acid was purchased from abcr GmbH. 14-pentadecenoic acid was purchased from Larodan AB. Maleimide-$PEG_8$-$CH_2CH_2COOH$ was purchased from PurePEG. All the other chemicals and solvents were purchased from Sigma-Aldrich.

**Synthesis of C6-6', C11-6', C11-3' and C14-6'**

**Step 1**: Heptakis-(6-deoxy-6-mercapto)-beta-Cyclodextrin (0.04 mmol, 50 mg) and bi-functional molecules bearing an allyl and a carboxylic acid end-groups (0.28 mmol, 36 mg 6-heptenoic acid, 55.5 mg 11-dodecenoic acid or 67 mg 14-pentadecenoic acid) were dissolved in 5 mL of DMSO. The reaction mixture was placed under ultraviolet (UV) lamp (250 W) and stirred overnight. The resulting cyclodextrins were precipitated by the addition of acetone (15 mL), collected by centrifugation and dried under vacuum.

**Step 2**: Cyclodextrin derivative obtained in step 1 (~ 0.02 mmol; 33 mg for C6-6', 38 mg for C11-6' or C11-3', 41 mg for C14-6'), N-hydroxysuccinimide (0.6 mmol, 57.5 mg), 1-Ethyl-3-(3-dimethylaminopropyl) carbodiimide (0.3 mmol, 57 mg) and 4-dimethylaminopyridine pyridine (0.01 mmol, 1.2 mg) were dissolved in 5 mL of DMSO and stirred overnight. The resulting cyclodextrins were sequentially washed with ice-cold $H_2O$ (20 mL), $CH_3CN$ (10 mL) and $Et_2O$ (10 mL), collected by centrifugation and dried under vacuum.

**Step 3**: Neu5Acα(2,6)-Galβ(1-4)-GlcNAc-β-ethylamine (5.6 μmol, 4 mg) and ~1.6 μmol of the cyclodextrin derivative obtained in the step 2 (3.3 mg for C6-6', 3.7 mg for C11-6', 3.9 mg for C14-6') dissolved in 1 mL DMSO. 15 μmol triethylamine (TEA) was added to reaction and the mixture was stirred overnight. The reaction product was diluted with 0.01 M phosphate buffer (pH: 7.5) and concentrated using amicon filters (MWCO: 3k). The resulting material was further washed with distilled water and lyophilized. In the case of C11'3, the reaction was conducted in the same conditions but with the -2,3 linked version of the trisachharide, Neu5Acα(2,3)-Galβ(1-4)-GlcNAc-β-ethylamine. The grafting of the trisaccharide onto the spacer-modified cyclodextrin was confirmed with $^1H$ and DOSY NMR studies (S13 to S15).

**Synthesis of P8-6'**

**Step 1**: Heptakis-(6-deoxy-6-mercapto)-beta-Cyclodextrin (0.04 mmol, 50 mg) was dissolved in 1 mL of DMSO and added to Maleimide-$PEG_8$-$CH_2CH_2COOH$ (0.28 mmol, 146 mg) dissolved in 5 mL of phosphate buffer (pH:6.8). The reaction mixture was placed under ultraviolet (UV) lamp (250 W) and stirred overnight. The modified β-cyclodextrin was purified with dialysis against Milli-Q $H_2O$ and lyophilized.

**Step 2**: Cyclodextrin derivative obtained in step 1 (~0.02 mmol, 63 mg), N-hydroxysuccinimide (0.6 mmol, 57.5 mg), 1-Ethyl-3-(3-dimethylaminopropyl) carbodiimide (0.3 mmol, 57 mg) and 4-dimethylaminopyridine pyridine (0.01 mmol, 1.2 mg) were dissolved in 5 mL of DMSO and stirred overnight. The resulting cyclodextrin was washed with DCM-$Et_2O$ mixture (20 mL, 3 times) collected by centrifugation and dried under vacuum.

**Step 3**: Neu5Acα(2,6)-Galβ(1-4)-GlcNAc-β-ethylamine (5.6 μmol, 4 mg) and the cyclodextrin derivative obtained in the step 2 ( ~1.6 μmol, 5.7 mg) were dissolved in 1 mL DMSO. 15 μmol triethylamine (TEA) was added to reaction and the mixture was stirred overnight. The reaction product was diluted with 0.01 M phosphate buffer (pH: 7.5) and concentrated using amicon filters (MWCO: 3k). The resulting cyclodextrin was washed with DCM-$Et_2O$ mixture (20 mL, 3 times) collected by centrifugation and dried under vacuum.

**Synthesis of C1-6'**

**Step 1**: Carboxymethyl-beta-Cyclodextrin sodium salt (0.04 mmol, 56 mg) N-hydroxysuccinimide(0.6mmol,57.5mg),1-Ethyl-3-(3-dimethylaminopropyl) carbodiimide (0.3 mmol, 57 mg) and 4-dimethylaminopyridine pyridine (0.01 mmol, 1.2 mg) were dissolved in 5 mL of DMSO and stirred overnight. The resulting cyclodextrin was sequentially washed with $CH_3CN$ (10 mL) and $Et_2O$ (10 mL), collected by centrifugation and dried under vacuum.

**Step 2**: Neu5Acα(2,6)-Galβ(1-4)-GlcNAc-β-ethylamine (5.6 μmol, 4 mg) and the cyclodextrin derivative obtained in the step 2 ( ~1.6 μmol, 3 mg) were dissolved in 1 mL DMSO. 15 μmol triethylamine (TEA) was added to reaction and the mixture was stirred overnight. The reaction product was diluted with 0.01 M phosphate buffer (pH: 7.5) and concentrated using amicon filters (MWCO: 3k). The resulting material was further washed with distilled water and lyophilized.

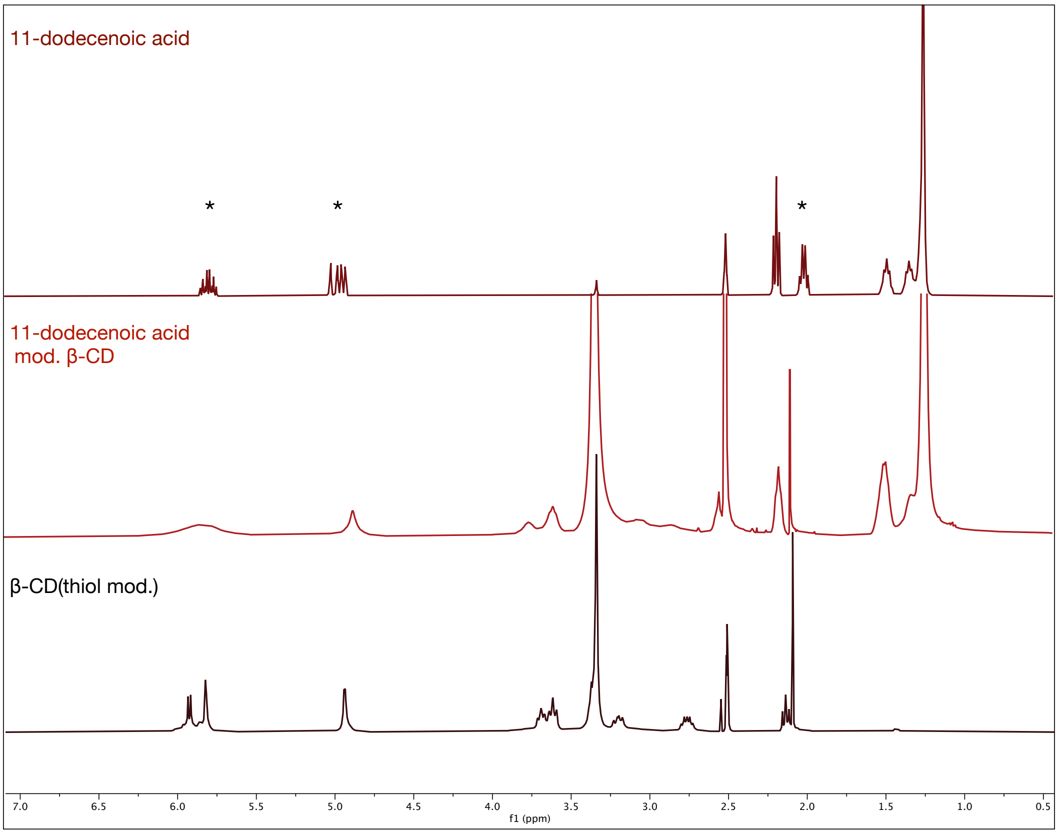


**Figure S11:** Stacked $^1$H NMR spectrum (in DMSO-d6) of 11-dodecenoic acid, 11-dodecenoic acid modified ß-clycodextrin and ß-clycodextrin (thiol modified). In the NMR of 11-dodecenoic acid, peaks from the hydrogens neighboring the double bond are demonstrated with *. These peaks move upfield upon binding to cyclodextrin.

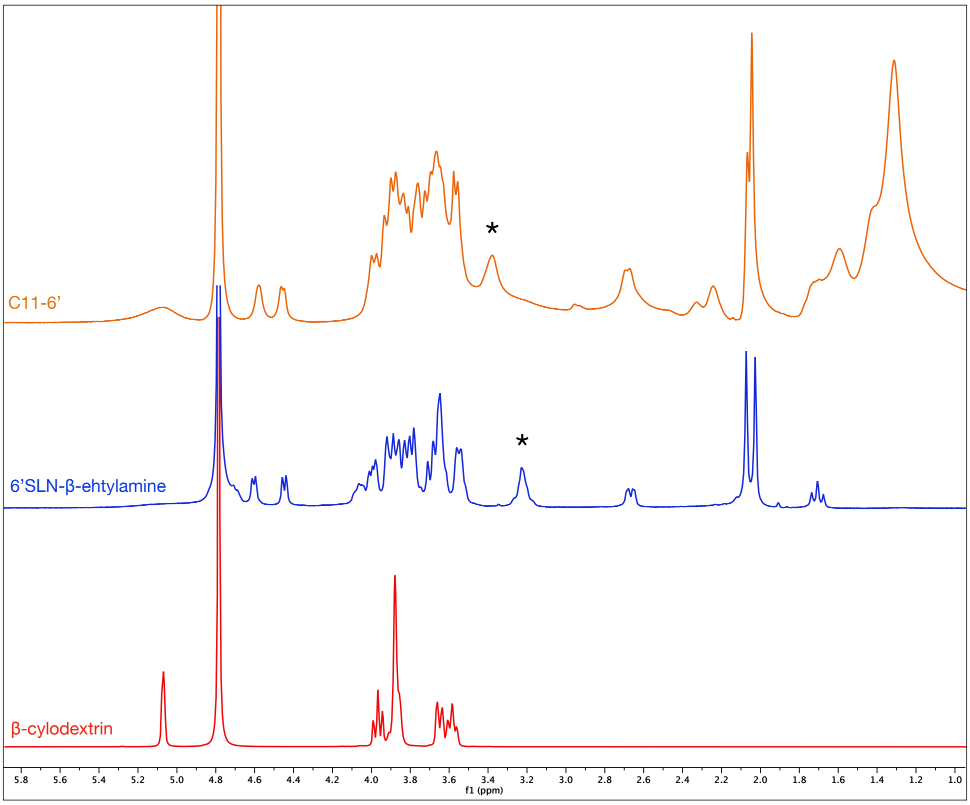


**Figure S11:** Stacked $^{1}$H NMR spectrum of ß-clycodextrin, 6’SLN-ß-ethylamine and C11-6’ in $D_2O$. The peak from the hydrogens neighboring the amine group (from 6’SLN-β-ehtylamine) is demonstrated with *. This peak moves slightly downfield upon binding to the linker.

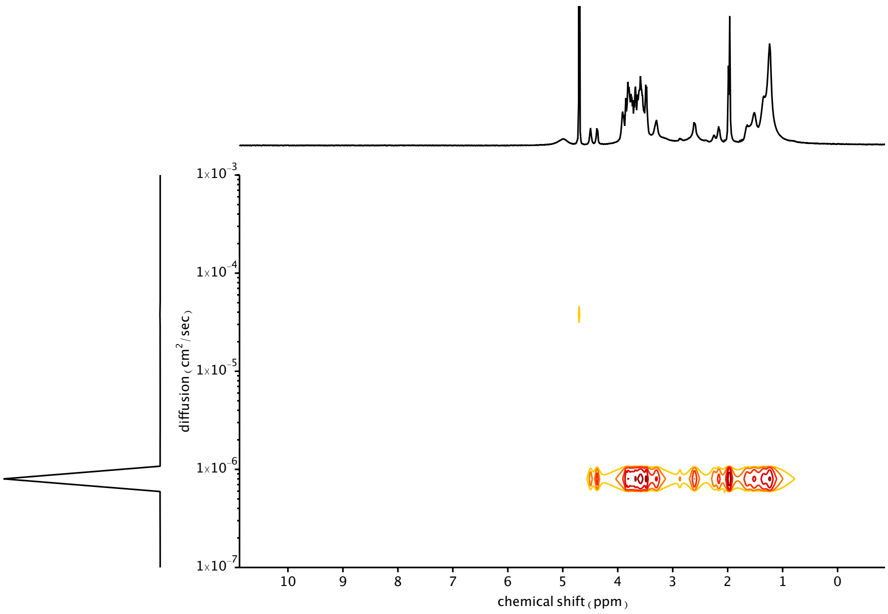


**Figure S13:** DOSY spectrum of the C11-6’ demonstrates that the resulting compound is free from unbound trisaccharides.

A

B

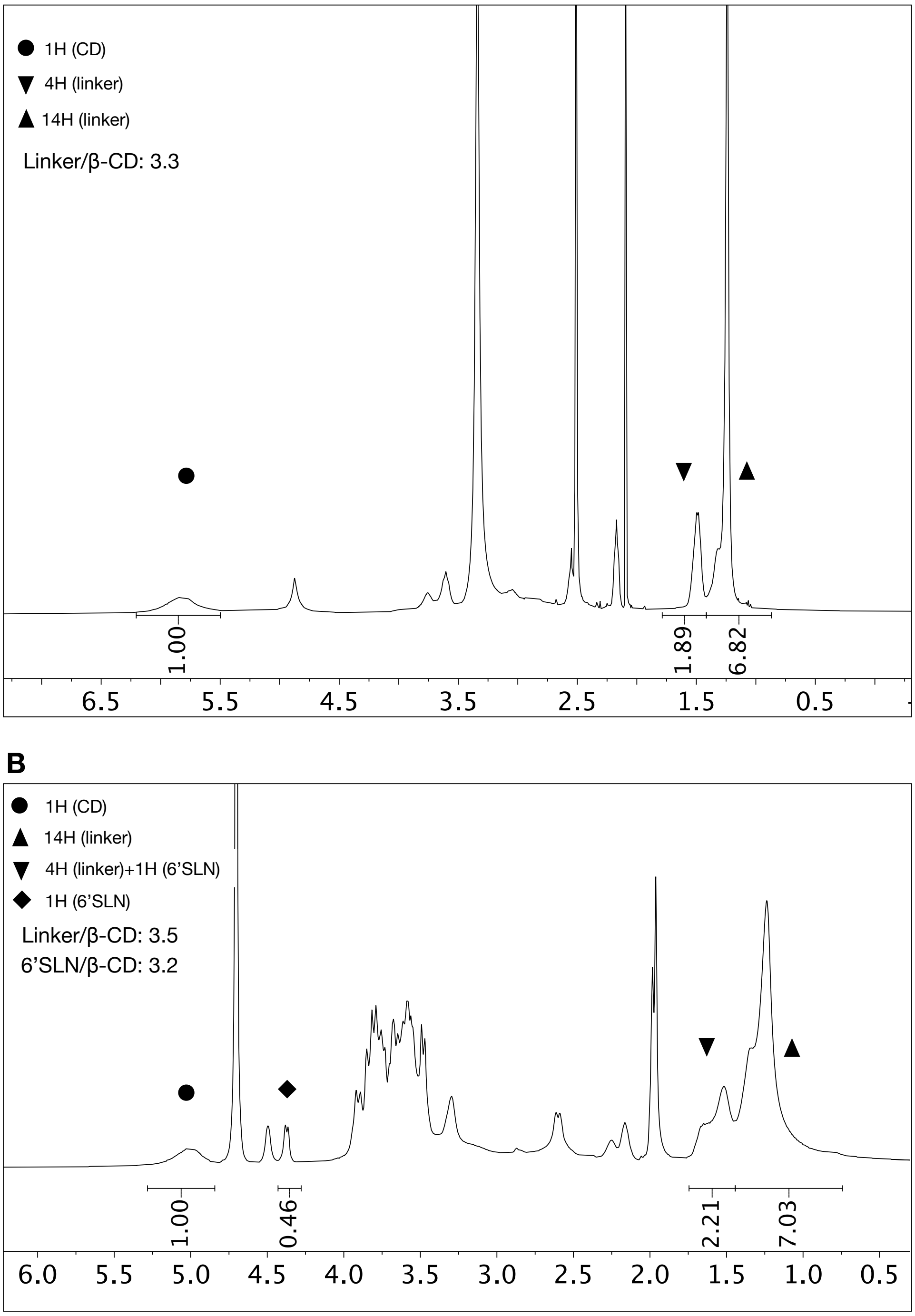


**Figure S14:** $^1$H NMR spectrum of intermediate of C11-6', cycodextrin modified with 11-dodecenoic acid, in DMSO-d6 (A) together with C11-6' (B).

### Biological Assays

**Materials:**

DMEM– Glutamax medium was purchased from Thermo Fischer Scientific. Tween 20® for washing buffer and 3,3'-diaminobenzidine (DAB) tablets were purchased from Sigma Aldrich. Primary antibody (Influenza A monoclonal antibody) was purchased from Light Diagnostics. Secondary antibody (Anti-mouse IgG, HRP-linked antibody) was purchased from Cell Signaling Technology®. The CellTiter 96® $AQ_{ueous}$ One Solution Cell Proliferation Assay that contains a tetrazolium compound [3-(4,5-dimethylthiazol-2-yl)-5-(3-carboxymethoxyphenyl)-2-(4-sulfophenyl)-2Htetrazolium, inner salt; MTS] and an electron coupling reagent (phenazine ethosulfate; PES) was purchased from Promega. Oseltamivir phosphate used in in vivo experiments was obtained from Roche (Palo Alto, CA) as a powder and prepared in sterile water for oral gavage (PO) administration of 0.1 ml.

**Cell Culture:**

MDCK (Madin-Darby Canine Kidney Cells) cell line, was purchased from ATCC (American Type Culture Collection, Rockville, MD). The cells were cultured in Dulbecco's modified Eagle's medium with glucose supplement (DMEM+ GlutaMAX™ ) containing 10% fetal bovine serum (FBS) and 1% penicillin/streptomycin. MDCK cell lines was grown in humidified atmosphere with $CO_2$ (5%) at 37°C.

**Viral Strains:**

A clinical isolate of HSV-2 was originally provided by Prof. M. Pistello, (University of Pisa, Italy) and was propagated and titrated by plaque assay on Vero cells. H1N1 Neth09 and B Yamagata were a kind gift from Prof M. Schmolke (University of Geneva). Avian strain NIBRG-23 (prepared by reverse genetics using A/turkey/Turkey/1/2005 H5N1 surface proteins and A/PR/8/34

(H1N1) backbone) was obtained from National Institute for Biological Standards and Controls, Potters Bar, United Kingdom and was grown further in 10 days old embryonated chicken eggs followed by virus purification and characterization. Clinical samples were provided from the Geneva University Hospital from anonymized patients. All influenza strains were propagated and titrated by ICC on MDCK cells in presence of TPCK-treated trypsin (0.2 mg/ml)

**Cell viability assay**

Cell viability was measured by the MTS [3-(4,5-dimethylthiazol-2-yl)-5-(3-carboxymethoxyphenyl)-2-(4-sulfophenyl)-2H-tetrazolium] assay. Confluent cell cultures seeded in 96-well plates were incubated with different concentrations of nanoparticles or ligand in triplicate under the same experimental conditions described for the antiviral assays. Cell viability was determined by the CellTiter 96 Proliferation Assay Kit (Promega, Madison, WI, USA) according to the manufacturer's instructions. Absorbance was measured using a Microplate Reader (Model 680, BIORAD) at 490 nm. The effect on cell viability at different concentrations of nanoparticles or cyclodextrins was expressed as a percentage, by comparing the absorbance of treated cells with the one of cells incubated with culture medium alone. The 50 % cytotoxic concentrations ($CC_{50}$) and 95 % confidence intervals (CIs) were determined using Prism software (Graph-Pad Software, San Diego, CA).

**Inhibition assays**

MDCK cells were pre-plated 24 h in advance in 96-well plates. Increasing concentrations of materials were incubated with the influenza virus (MOI: 0.02 or 0.01 for H5N1 and 0.1 for other viruses) at 37 °C for one hour and then the mixtures were added to cells. Following the virus adsorption (1 h at 37 °C), the virus inoculum was removed, the cells were washed and the fresh medium was added. After 24 h of incubation at 37°C, the infection was analyzed with immunocytochemical (ICC) assay.

The cells were fixed and permeabilized with methanol. Then the primary antibody (1:100 dilution) was added and incubated for 1 hour at 37°C. The cells were washed with wash buffer (DPBS + Tween 0.05%) three times; then secondary antibody (1:750 dilution) was added. After 1 hour the cells were washed and the DAB solution was added. Infected cells were counted and percentages of infection were calculated comparing the number of infected cells in treated and untreated conditions.

In the case of H5N1, flow cytometry-based inhibition assays were conducted additionally. MDCK cells were pre-plated 24 h advanced in a 24- well plate (75,000 cells/well). Increasing concentrations of materials were incubated with the influenza virus (MOI: 0.04) at 37 °C for one hour and then the mixtures were added to cells. Following the virus adsorption (1 h at 37 °C), the virus inoculum was removed, the cells were washed and the fresh medium was added. After 5 h of incubation at 37°C, the infection was analyzed with flow cytometry. Briefly cells were trypsinized and fixed using IC fixation buffer (Thermo Fisher Scientific, Netherlands) for 15 minutes at room temperature followed by permeabilization using 1X IC permeabilization buffer (Thermo Fisher Scientific, Netherlands) for 15 minutes at 4°C. Cells were stained with Anti-Influenza A Virus Nucleoprotein mouse monoclonal antibody [D67J] (FITC) (Abcam ab210526, The Netherlands) for 30 minutes at 4°C. Antibody dilution was 1:80 in permeabilization buffer and 50 $\mu$l per tube was added. FACS analysis was carried out using FACS Calibur 3 software. The concentration producing 50 % reduction in the number of infected cells (effective concentration ($EC_{50}$)) was determined using the Prism software.

H7N1 infectivity was evaluated through Luciferase activity. MDCK cells were seeded at $5x10^4$ on 96-wells plates. After 24h, the medium was replaced by serum-free medium. Increasing concentrations of C11-3' were incubated with 100 pfu of H7N1 A/Turkey/Italy/977/1999 encoding the NanoLuciferase. The mixture was incubated 1h at 37°C before to be added to the cells for another 1h incubation at 37°C (100 µL per well). Cells were washed and medium replaced by serum-free medium with 1 µg/mL TPCK-Trypsin. Twenty-four hours post-infection, cells were washed and then lysed with 40 µl per well of Nano Glo® Luciferase Assay Buffer (Promega) diluted 1/2 in PBS. Luciferase activity was measured in the cell lysates using a Tecan Infinite M200PRO plate reader: 15 µl of Nano Glo® Luciferase Assay Substrate (Promega) diluted 1/5000 in PBS were added to 15 µl of lysate for each well.

### Virucidal Assays

Viruses (focus forming unit (ffu):$10^5$/mL) and the materials ($EC_{99}$ concentration, figure S12) were incubated for 1 hour at 37°C. Serial dilutions of the virus-material complex together with the non-treated control were conducted and transferred onto the cells. After 1 hour, the mixture was removed and the fresh medium was added.  Next day, viral titers were evaluated with ICC assay. For the ICC assay, the same procedure described above was followed.

**Figure S12:** The material concentrations at which the in-vitro virucidal assays were performed.

| | Material | Concentration (µg/mL) |
|---|---|---|
| A/Netherlands/2009 H1N1 | C6-6’ | 100 |
| | C11-6’ | 100 |
| | C14-6’ | 100 |
| | P8-6’ | 500 |
| | C11-3’ | 500 |
| | NPs C15-6’SLN | 100 |
| | NPs PEG4-6’SLN | 500 |
| A/Singapore/2004 H3N2 | C11-6’ | 100 |
| B/Wisconsin/2010 | C11-6’ | 200 |

### Data analysis

All results are presented as the mean values from three independent experiments performed in duplicate. The $EC_{50}$ values for inhibition curves were calculated by regression analysis using the program GraphPad Prism version 5.0 (GraphPad Software, San Diego, California, U.S.A.) to fit a variable slope-sigmoidal dose–response curve.

### Ex vivo

Co-treatment: Human airway epithelia reconstructed *in vitro,* MucilAir tissues, (Epithelix Sàrl, Geneva, Switzerland) were cultured at the air-liquid interface from a mixture of nasal polyp epithelial cells originating from healthy donors. Influenza H1N1 pdm 2009 clinical strain (1e4 rna copies/tissue) and C11-6' (50 µg/tissue) were transferred onto tissues without pre-incubation, together with non-treated control. After 4 hours of incubation time, the tissues were washed twice. On a daily basis, the basal medium was changed. To conduct daily qPCR measurements, 200 uL of medium was added onto tissues and then collected 20 minutes later. RNA extracted with EZNA viral extraction kit (Omega Biotek) was quantified by using qPCR with the QuantiTect kit (#204443; Qiagen, Hilden, Germany) in a StepOne ABI Thermocycler.

Post treatment: MucilAir tissues were infected with H1N1 pdm 2009 (1e4 copies/tissue) after 4 h the inoculum was removed and tissues were washed. After 20 h an apical wash was done for 20' and subsequently C11-6' (30 µg/tissue), or an equal volume of medium in the untreated tissues, were added apically. Everyday after the 20' apical wash new addition of C11-6' was performed. RNA was then extracted and qPCR done as described above. A similar procedure in the absence of virus was conducted for toxicity studies.

#### Immunofluorescence

Influenza infected cells were detected by direct with Influenza A antibody (Light Diagnostic) and beta tubulin primary rabbit antibody (Abcam) was used as a marker of ciliated cells. The Alexa 488-

goat anti-rabbit Ab and the Alexa 594-goat anti-mouse Ab (Life technologies) were used as secondary Ab and nuclei were stained with DAPI. Images were acquired with Zeiss LSM 700 Meta confocal microscope and processed by Imaris.

**Lactate dehydrogenase assay (LDH)**

LDH release in basal medium was measured with the Cytotoxicity Detection Kit (Roche 04744926001).

**MTT assay**

MTT solution was diluted in MucilAir medium (1 mg/ml) and 300 μl were added basally in a 24 well plate. After 4 hours incubation at 37°C the tissues were transferred in a new plate and lysed with 1 ml of DMSO. The supernatant was read at 570 nm. Percentages of viability were calculated comparing treated and untreated tissues.

**ELISA**

Interleukin-6 (IL-6), CXC motif chemokine 10 (CXCL10 or IP-10), CC motif chemokine5 (CCL5 or RANTES), interleukin-8 (IL-8 or CXCL-8) and interferon lambda (IL-29 /IL-28B) were measured in the basal medium by ELISA (R&D DY206-05, DY266-05, DY278-05, DY208-05 and DY1598B-05) following everyday treatment with different concentrations of CD.

**In vivo**

**Single treatment**

Two groups of five BALB/c mice were treated at day 0 with 50μl of PBS or C11-6' (1.25 mg/kg) and immediately inoculated with A/NL/09 ($10^2$ ffu). Two days post-infection mice were euthanized. Lung homogenate and bronchoalveolar lavage were collected to quantify the viral titer through titration or qPCR measurements. The C11-6' treated group was retreated with the same amount of the compound. After the tissue disruption, the RNA was extracted with Trizol and quantified by using qPCR while BAL were subjected to plaque assay. Two independent experiments were performed.

**Co and post-treatment**

Female 20 g BALB/c mice were obtained from Charles River Laboratories (Wilmington, MA) for this investigation. The animals were quarantined for 6 days prior to use and maintained on standard rodent chow and tap water ad libitum. Mouse adapted influenza A/California/04/2009 (H1N1pdm) was kindly provided by Dr. Elena Govorkova (St. Jude Children's Research Hospital, Memphis, TN). The virus stock was prepared from an infection in Madin-Darby canine kidney (MDCK) cells and administered to the mice intranasally (IN) in 75 μl (approximately 3 CCID50).

Pre treatment: mice (12 per experimental treatment group, and 10 for positive control group) were anesthetized by IP injection of ketamine/xylazine (50/5 mg/kg) followed by 6'SLN CD treatments of 3.75, 1.25 mg/mg and the vehicle placebo (water) administered in a 0.05 ml volume by IN about 10 minutes prior to the IN infection of a 75 μl suspension of influenza virus inoculum containing 3 $CCID_{50}$ challenge dose of virus. Oseltamivir (30 mg/kg/day) treatments were administered with per os, twice daily for 5 days starting 1h before infection as the positive control. Individual weights were recorded everyday beginning of the day of virus challenge and mice were observed daily for survival.

Post treatment: mice (n=10 per group) were anesthetized by IP injection of ketamine/xylazine (50/5 mg/kg) daily treatments of 7.5, or 3.75, and the vehicle placebo (PSS) administered in a 0.05 ml volume by intranasal beginning 8 h or 24 h post-challenge with influenza virus. Oseltamivir (30 mg/kg/day) treatments were administered per os, twice daily (bid) for 5 days starting at 8 h or 24 h post-infection as the positive control. The oseltamivir-treated mice also received a 0.05 mL PSS placebo by IN once daily to mimic the 6'SLN CD-treatment conditions. Individual weights were recorded every day beginning on the day of virus challenge and mice were observed daily for survival. Survival curves were compared with the Mantel-Cox log-rank test.

**Ethics statement**

This study was carried out in accordance with INRA guidelines in compliance with European animal welfare regulation. The protocols were approved by the Animal Care and Use Committee at "Centre de Recherche de Jouy-en-Josas" (COMETHEA) under relevant institutional authorization ("Ministère de l'éducation nationale, de l'enseignement supérieur et de la recherche"), authorization number: 2015100910396112v1 (APAFIS#1487). All experimental procedures were performed in a Biosafety level 2 facility.

This study was conducted in accordance with the approval of the Institutional Animal Care and Use Committee of Utah State University dated September 25, 2019 (expires Nov. 28, 2021). The work was done in the AAALACaccredited Laboratory Animal Research Center of Utah State University.
